# Engineering nanoparticles that target fibroblast activation protein in cardiac fibrosis

**DOI:** 10.1101/2025.04.23.650330

**Authors:** Chima V. Maduka, Gabriel J. Rodriguez-Rivera, Manuela Garay-Sarmiento, Amy R. Perry, Sara Boyd, Kendra Worthington, Mackenzie Obenreder, Stella Rydahl-Kim, Francis G. Spinale, Jason A. Burdick

## Abstract

Cardiac fibrosis and dysfunction, hallmarks of debilitating heart disease, are driven by fibroblast activation protein alpha (FAP). The specificity of FAP to the disease creates an opportunity for directed drug delivery, as FAP delineates the fibrotic region. We leverage this vulnerability to target the fibrotic region of the heart, using anti-FAP antibody-modified nanoparticles (NPs) that encapsulate and release the highly specific FAP inhibitor, talabostat. NP crosslinking with an FAP-sensitive peptide attenuates passive release, which is then accelerated in the presence of FAP. Intravenous administration of these NPs results in reversal of established cardiac fibrosis and dysfunction in a rat model of myocardial infarction, using a talabostat dose that is 400,000-fold lower than that used in clinical trials. This innovative and clinically translatable strategy enables targeted drug delivery, overcoming limitations of systemic approaches by reducing therapeutic dose, minimizing off-target effects, and accommodating patient-specific variability in FAP expression.

## Introduction

Fibrosis is a hallmark of organ dysfunction in numerous conditions, including heart failure (HF), where cardiac dysfunction triggers neurohormonal systems that are initially compensatory, but become maladaptive^1^. Clinical therapeutics for HF, including drugs that target neurohormonal systems like the renin–angiotensin system, the mineralocorticoid receptor, the sympathetic nervous system and the natriuretic system reduce mortality^2^, but fail to address the molecular progression of the disease and the heterogeneity of clinically encountered HF patients^3^. Thus, current therapies are intrinsically non-curative^1^. To make matters worse, HF disproportionately affects Black and Hispanic individuals, who have the highest incidence and prevalence of HF, with earlier onset, higher hospitalization costs and increased death rates^4,5^.

Among patients presenting with HF arising from several etiologies such as myocardial infarction (MI)^6^, hypertrophic cardiomyopathy and dilated cardiomyopathy^7^, fibroblast activation protein alpha (FAP) has emerged as a key deleterious enzyme that molecularly orchestrates extensive cardiac fibrosis. Initially triggered by interleukin-1β signaling from immune cells^8^, FAP is a potent inflammatory driver^9^ that fosters cardiac dysfunction and aberrant extracellular matrix remodeling by targeting fibrillar (type 1) collagen^10^. In fact, FAP in the circulation^11^ and imaging of FAP in the diseased heart via positron emission tomography^12^ are actively being explored as biomarkers to identify patient subsets who will benefit from therapies tailored to FAP for precision medicine^13^.

Yet, substantial hurdles exist toward realizing inclusive HF therapies against FAP. For instance, FAP inhibition through the systemic administration of small molecule inhibitors produces only marginal benefits due to insufficient drug localization in the diseased heart^14^, with escalating drug doses causing off-target toxicities^15,16^. Chimeric antigen receptor (CAR) T-cell therapy against FAP has shown immense promise in reversing established cardiac fibrosis and dysfunction in HF^7,17^ and is poised to positively impact certain patient subsets. However, it is limited in availability^18^ and by the patient health status (e.g. malnutrition)^19^, neurotoxicity and infection^20,21^ and life-threatening adverse effects, such as cytokine release syndrome^21^, which disproportionately affects older^20^ and Hispanic^22^ populations. Thus, new FAP-targeted therapies are needed that overcome these limitations.

## Results and Discussion

To address the need for new inclusive technologies to treat HF patients, we engineered a nanoparticle (NP) platform that (i) targets the diseased region of the heart via (anti-) FAP antibodies and (ii) releases an inhibitor of FAP to alter and reverse disease progression. Specifically, we demonstrate that NPs can encapsulate the highly specific FAP inhibitor, talabostat, through the charge-driven interaction between acrylated hyaluronic acid (HA) and chitosan (Fig. 1a). Talabostat has been advanced to Phase II cancer clinical trials^15,16^ and is historically ascribed to inhibit FAP enzymatic activity^23^. We extend this observation by showing that talabostat reduces FAP expression in cultured, activated human cardiac fibroblasts (Supplementary Fig. 1a-e), likely by the internalization of the serine protease receptor-inhibitor complex^24^, similar to observations made in systemic sclerosis^25^.

**Figure 1.**
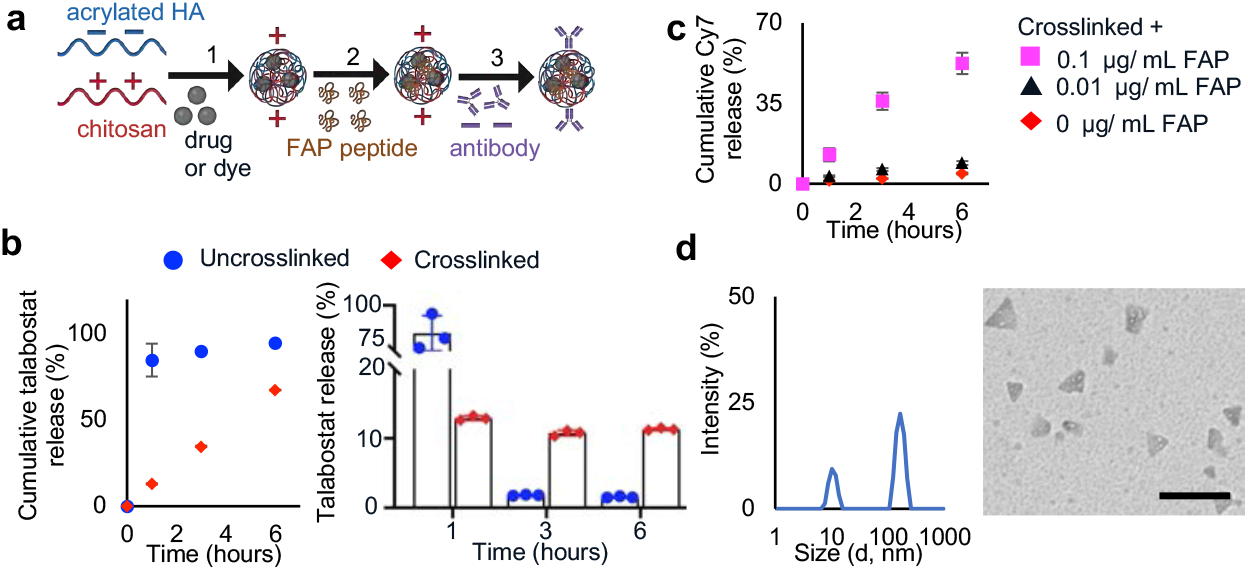
Engineered nanoparticles (NPs) that bind and are responsive to fibroblast activation protein alpha (FAP). **a**, Charge-driven interaction between solutions of chitosan (containing talabostat or Cyanine 7, Cy7) and acrylated hyaluronic acid (HA) form NPs (step 1) that can be crosslinked by thiolated FAP peptides via Michael Addition (step 2); negatively charged (anti-FAP or IgG1) antibodies are then adsorbed to positively charged NPs (step 3). Intravenous administration of NPs with adsorbed anti-FAP antibodies allows targeting to the fibrotic region of the diseased heart. **b**, Cumulative (left, reported as amount released between 0-1 hours, 1-3 hours, 3-6 hours) and time-specific release (right) of talabostat (quantified through activity against FAP) from both uncrosslinked and crosslinked NPs. **c**, Cy7 release from crosslinked NPs with and without recombinant human FAP enzyme. **d**, Size distribution and transmission electron microscopy images (scale bar, 100 nm) of crosslinked NPs adsorbed to anti-FAP antibodies. Mean (SD), n = 3 biological replicates.

HA and chitosan have been previously self-assembled as NPs^26,27^; here, we introduce a modified HA that allows NP crosslinking to attenuate the rapid, initial release of encapsulated talabostat prior to targeting the heart^28,29^. Specifically, the acrylate groups in modified HA (Supplementary Fig. 2) allowed the crosslinking of NPs by thiolated, FAP-cleavable peptides via a Michael Addition reaction to limit early release (Fig. 1b), overcoming a major hurdle in clinical translation^28,29^. Direct mass spectrometry analysis of released talabostat proved unfeasible due to its cyclization and challenging ionization properties^30^. Consequently, we adopted an indirect fluorogenic enzymatic assay that utilizes recombinant human FAP enzyme to profile talabostat release (Fig. 1b)^15^. Uncrosslinked NPs showed a burst release of nearly 85% of talabostat, whereas the crosslinked NPs showed a gradual release of talabostat over time.

To illustrate the responsiveness of crosslinked NPs to FAP, we incorporated cyanine 7 (Cy7) dye within NPs. The incorporation of Cy7 was necessary because profiling talabostat itself inherently requires FAP addition, precluding its use as an independent measure of NP responsiveness. Moreover, the incorporation of Cy7 facilitates visualization of NP distribution within tissues during subsequent in-vivo studies. Validating our findings with talabostat, crosslinking NPs limited the initial burst release of Cy7 (Supplementary Fig. 3a). However, we observed slower release of Cy7 compared with talabostat, likely attributable to the dye’s 2.4-fold higher molecular weight (Supplementary Fig. 3a, Fig. 1b). Additionally, crosslinked NPs exposed to different FAP enzyme concentrations (to simulate varied FAP expression observed in patients^31^) released Cy7 in a dose-dependent manner, demonstrating enzyme-responsivity (Fig. 1c).

NPs were engineered to have a net positive charge, such that negatively-charged, monoclonal antibodies are adsorbed onto NPs, resulting in a dose-dependent decrease in NP charge (Supplementary Fig. 3b-c). Beyond the NP to antibody (weight by weight) ratio of 1:9, there was no significant change in NP charge with the introduction of additional FAP (FAP NPs) or IgG1 (IgG NPs) antibodies (Supplementary Fig. 3b-c); thus, a 1:9 ratio was deemed optimal for antibody modification and was used for subsequent experiments. Compared to NPs alone, NPs with adsorbed monoclonal antibodies had similar mean hydrodynamic diameters and polydispersity indices (Supplementary Fig. 3d-e). Size distribution data showed peaks consistent with NPs, as well as peaks at approximately 10 nm, which may indicate the presence of unadsorbed, excess antibodies (Supplementary Fig. 3f; Fig. 1d). Furthermore, dried NPs, with and without adsorbed antibodies, were irregularly shaped on transmission electron microscopy (Supplementary Fig. 3g; Fig. 1d). To determine whether FAP NPs are preferentially taken up by FAP-expressing fibroblasts over IgG NPs, human activated, primary, cardiac fibroblasts were exposed to fluorescein isothiocyanate-dextran (FITC-dextran)-containing NPs^32^. We observed preferential cellular uptake of FAP NPs in comparison to IgG NPs, showing specificity in targeting (Supplementary Fig. 4a-b).

Localization of NPs to diseased tissue is essential for their therapeutic effect. To investigate this, Cy7-containing NPs adsorbed with either FAP or IgG antibodies were intravenously administered to a clinically-relevant ischemia-reperfusion rat model of MI (ligation of the left anterior descending (LAD) artery for 30 minutes followed by reperfusion)^33^. Specifically, NPs were administered 3 or 7 days after MI or in healthy rats, and organs were harvested one hour later and imaged (Fig. 2a). These time points (days 3 and 7 post-MI) were carefully selected as they have been shown to coincide with extensive cardiac fibrosis^34^, markedly elevated FAP expression^6,11,34^ and cardiac dysfunction measured by echocardiography^35^, metrics indicative of HF^7,17^, which we validated ourselves (Supplementary Fig. 5a-d). Moreover, in human patients, FAP levels peak from day 3 post-MI^11^, making the selected injection time points biologically and clinically relevant, as it follows the acute (48 hr) period after MI that is often associated with complications^36^ and coincides with potential hospital discharge post-MI.

**Figure 2.**
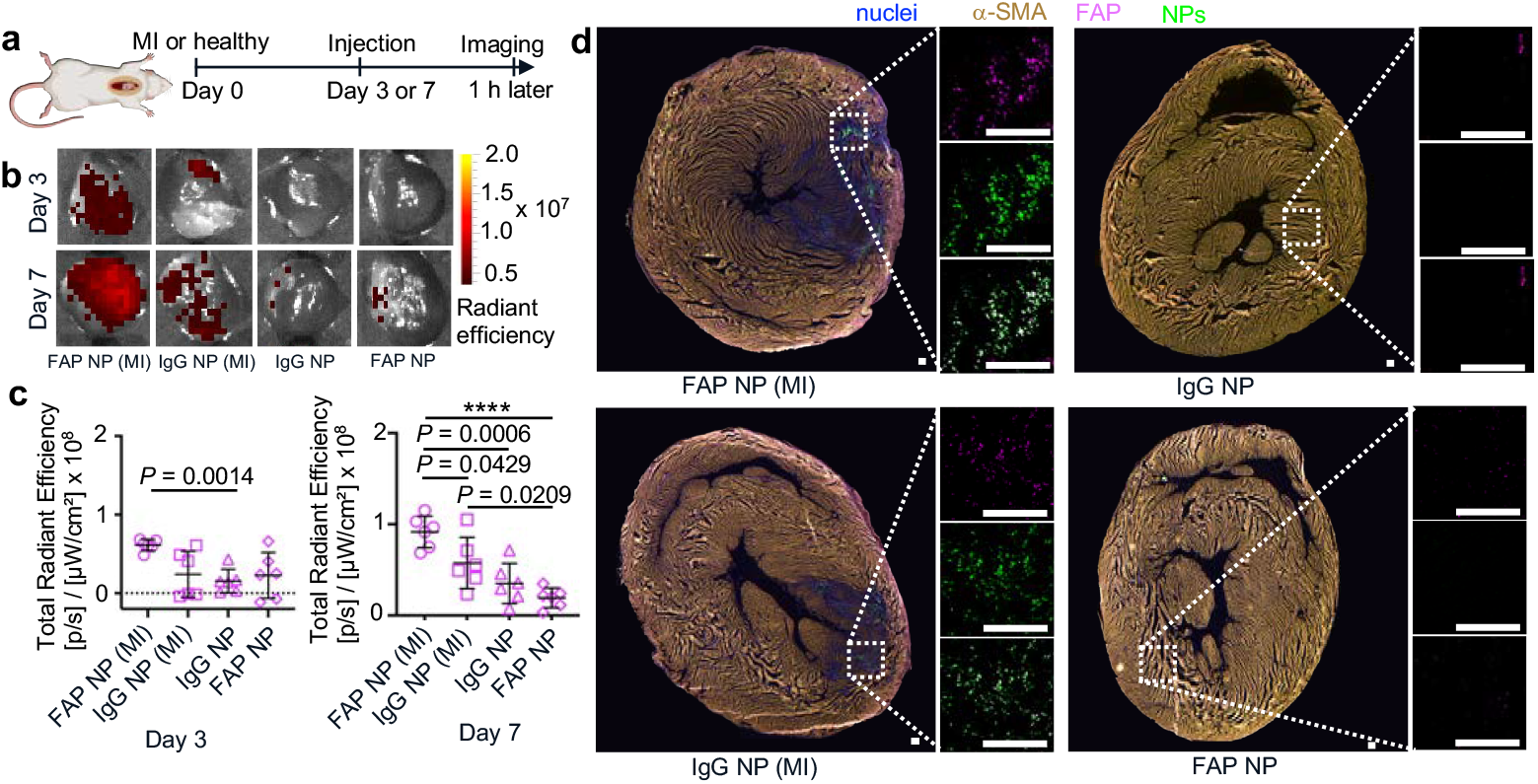
Administered NPs target the diseased region of the heart. **a**, Alongside healthy controls, rats were intravenously administered NPs (containing Cy7) with adsorbed FAP (FAP NP) or IgG (IgG NP) antibodies, 3 or 7 days following myocardial infarction (MI). One hour post-injection, organs were collected for ex-vivo analyses via in vivo imaging system (IVIS) and immunohistochemistry (IHC). **b**, Whole heart images and **c**, quantification of epifluorescence from IVIS images. **d**, IHC images (with magnification) of axial heart sections from Day 7 treatments; scale bar, 200 μm. Mean (SD); Brown-Forsythe and Welch ANOVA followed by Dunnett’s T3 multiple comparisons test (c, day 3) or one-way ANOVA followed by Tukey’s multiple comparison test (c, day 7); n = 6 rats per group, **** *P* < 0.0001.

Upon imaging one hour after administration at either 3 or 7 days, there was little NP signal observed in healthy animals (Fig. 2b-c; Supplementary Fig. 6), although NP signals were observed in the kidneys, spleen, lungs and liver (Supplementary Fig. 6), whose natural clearance of NPs has been exploited to treat fibrosis in these organs^37–39^. However, in comparison to other groups, FAP NPs were observed to target MI hearts, with significantly enhanced targeting at 7 days compared to 3 days post-MI (Fig. 2b-c; Supplementary Fig. 6), consistent with the higher FAP expression at this later time point^6^. This was validated by immunohistochemistry (IHC), which shows NPs localized to fibrotic regions of the heart after delivery at day 3 (Supplementary Fig. 7) and day 7 (Fig. 2d) after MI, coincident with FAP staining, underscoring the ability of engineered NPs to target fibrosis by penetrating the remodeling extracellular matrix^40^. Additionally, IHC confirmed the near absence of NPs in healthy controls as well as lower amounts of IgG NPs that still accumulate in the diseased heart (Fig. 2d; Supplementary Fig. 7) likely due to enhanced permeability and retention during inflammation^41^, traditionally relied upon for NP delivery immediately after MI^33,42^.

To assess the influence of talabostat delivery on cardiac function and fibrosis, NPs containing talabostat were administered intravenously on days 3 and 7 after MI and then analyzed via echocardiography and histology on day 30 after MI (Fig. 3a). Sham controls (open thoracotomy but without ligation of the LAD), MI groups that received no treatment, intravenous administration of the equivalent amount of talabostat in normal saline, and NPs without talabostat (empty NPs) were included for comparison. As expected, MI resulted in decreased fractional shortening, decreased ejection fraction, and increased left ventricular internal diameter at end systole (LVIDs) relative to sham controls (Fig. 3b-d). In comparison to MI groups, systemically administered talabostat in normal saline and empty NPs marginally increased fractional shortening and ejection fraction and reduced LVIDs, but did not restore these metrics of cardiac function to levels of the sham controls (Fig. 3b-d). Marginal therapeutic effects with empty NPs may be due to antibody-receptor complex formation, triggering a phagocytotic response. Remarkably, NPs encapsulating talabostat not only restored fractional shortening, ejection fraction, and LVIDs to levels similar to sham controls, likely by localizing delivery, but they showed efficacy in fractional shortening over systemically administered talabostat and empty NPs (Fig. 3b-d). The capability of NPs encapsulating talabostat to normalize cardiac function was consistent across left ventricular internal diameter at end diastole, end-systolic and end-diastolic volumes, with heart rates unchanged across groups (Supplementary Fig. 8a-d).

**Figure 3.**
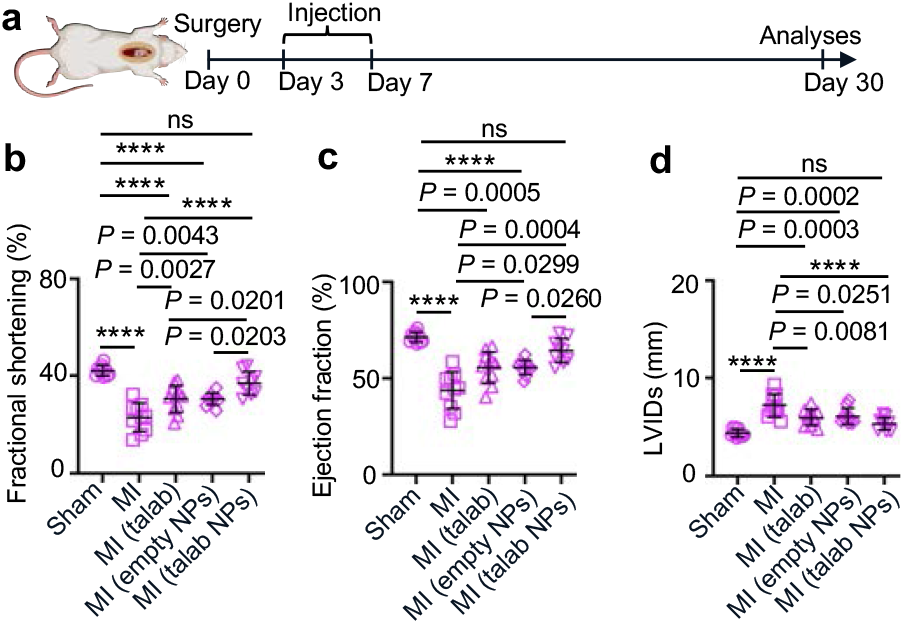
FAP-inhibitor encapsulated NPs restore cardiac function. **a**, Rats were injected 3 and 7 days following myocardial infarction (MI) or sham surgery, with subsequent analyses at day 30. **b-d**, Cardiac function was assessed by fractional shortening (b), ejection fraction (c) and left ventricular internal diameter at end systole (LVIDs; d) for sham controls, MI alone, and MI with intravenous talabostat in saline (talab), empty NPs or NPs encapsulating talabostat (talab NPs). Mean (SD); one-way ANOVA followed by Tukey’s multiple comparison test (b,d) or Brown-Forsythe and Welch ANOVA followed by Dunnett’s T3 multiple comparisons test (c); n = 10 rats per group (sham, MI, empty NPs), n = 11 (talab), n = 9 (talab NPs); not significant (ns), **** *P* < 0.0001.

In this study, each rat received a dose of 0.125 ng of talabostat via NPs at each injection time point, which is 27,000-fold lower than the half-maximal inhibitory concentration (IC50) of talabostat (173. 7 ng/ mL), assuming each rat weighing 300 g has 20 mL of blood^43^. For context, during cancer clinical trials, 200 mcg of talabostat was administered twice daily to patients for 6 cycles (each cycle consisted of 14 consecutive days) to maintain therapeutic levels in circulation, and this dose had to be lowered (thereby compromising efficacy) after many patients exhibited adverse events^16^. Relative to tested doses and in patients weighing 100 kg, our NP platform is projected to normalize cardiac function with a 400,000-fold reduction in clinical trial doses. This represents a major advancement in targeted drug delivery to the diseased heart, enabling a substantial reduction in therapeutic doses, thus minimizing associated toxicities. This progress is particularly exciting considering the clinical relevance of polymeric NPs, as evidenced by the numerous formulations already approved by the U.S. Food and Drug Administration (FDA) or currently undergoing clinical trials^44^.

To understand the process by which our engineered NP platform restores cardiac function, we histologically assessed infarct area by 2,3,5-triphenyltetrazolium chloride (TTC) staining and observed that NPs encapsulating talabostat significantly reduced left ventricular infarct area compared with MI groups that received no treatment, systemically administered talabostat or empty NPs (Supplementary Fig. 9a-c). Furthermore, in contrast to other groups, only NP-mediated delivery of talabostat reduced FAP expression to levels comparable to sham controls (Fig. 4a; Supplementary Fig. 9d), likely by the internalization of the serine protease receptor-talabostat complex^24^ and consistent with our observation in cultured human cardiac fibroblasts (Supplementary Fig. 1a-e). Given the cause-effect relationship between FAP expression and cardiac fibrosis^14^, we next evaluated metrics of cardiac fibrosis. We observed that NPs encapsulating talabostat significantly reduced the fibrotic region of the left ventricles when compared with MI groups that received no treatment, systemically administered talabostat, or empty NPs (Fig. 4b-c); this observation was corroborated by Picrosirius red staining under regular and polarized light microscopy (Supplementary Fig. 9e).

**Figure 4.**
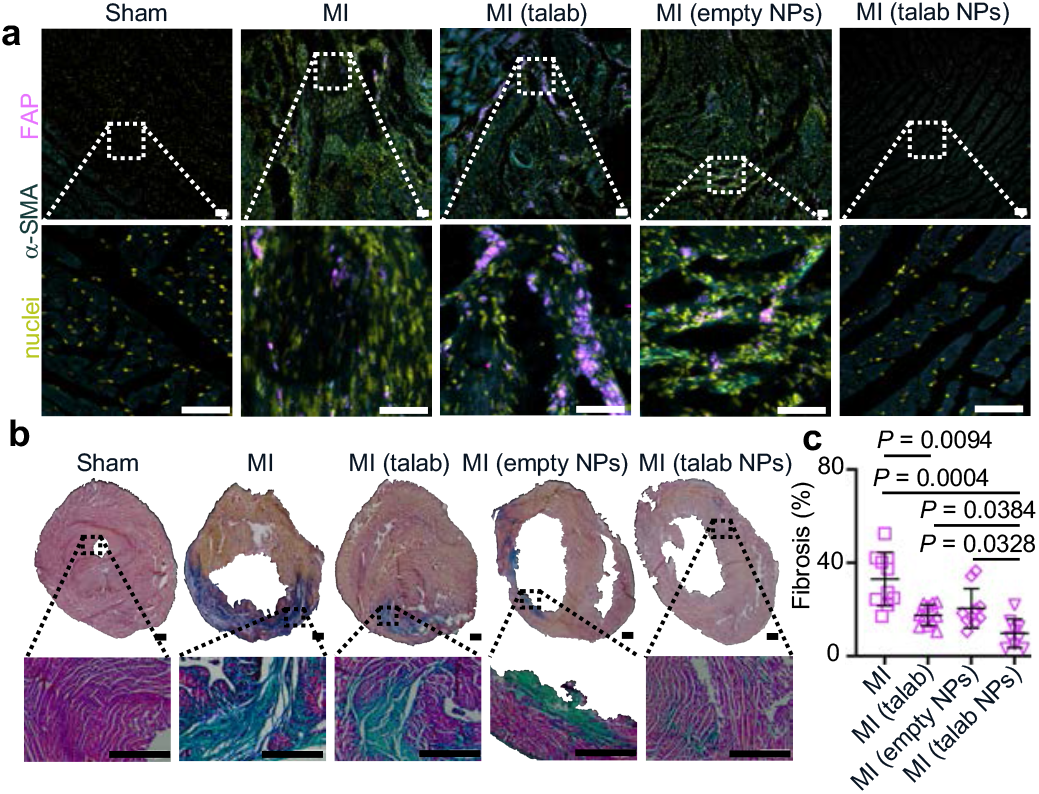
FAP-inhibitor encapsulated NPs reduce cardiac fibrosis. **a**, Immunohistochemical images (with magnification) of the left ventricle (LV) of rats that underwent sham surgery, MI alone, and MI with intravenous talabostat in saline (talab), empty NPs, or NPs encapsulating talabostat (talab NPs); scale bar 50 μm. **b**, Masson Trichrome staining and **c**, quantification of LV fibrosis of heart axial sections (with magnification) for the various groups; scale bar 1 mm. Mean (SD); Brown-Forsythe and Welch ANOVA followed by Dunnett’s T3 multiple comparisons test; n = 10 rats per group (sham, MI, empty NPs), n = 11 (talab), n = 9 (talab NPs).

During clinical trials with talabostat, key indices of toxicity were anorexia (leading to weight loss) and peripheral edema^15,16^, both of which were not observed in treated rats in our study. Experimental induction of MI resulted in weight loss compared to sham controls (Supplementary Fig. 10a); however, compared to MI groups that received no treatment, rats that received NPs encapsulating talabostat showed significantly improved weight gain (Supplementary Fig. 10a). Additionally, compared to untreated groups, the administration of NPs encapsulating talabostat did not result in observable histopathological changes (Supplementary Fig. 10b) in the liver, kidney, lungs, and spleen, where NPs were biodistributed, or in the skin, where FAP may be expressed during wound healing^17^.

In summary, cardiac fibrosis remains a major clinical challenge. Yet, it paradoxically presents a unique opportunity for the targeting of therapeutics based on hallmark fibrotic signatures, such as FAP, which define affected areas. Prior biomaterial strategies have largely focused on limiting the progression of cardiac fibrosis and dysfunction, including through targeting proteases^14,33,45–47^. In contrast, our approach reverses established cardiac fibrosis and dysfunction via the engineering of a NP platform that not only targets FAP-related regions of the tissue, but delivers a therapeutic FAP inhibitor. While similar in approach and therapeutic outcome, our strategy obviates substantial issues associated with CAR T-cell therapies^18–22^ for cardiac fibrosis, creating alternative technologies for underserved patient populations to advance inclusive healthcare. Moreover, our method overcomes the limitations of systemic drug administration by locally releasing drugs, an approach that markedly lowers the therapeutic dose needed and limits off-target effects. By releasing drugs in a manner directly proportional to FAP levels, our nanotechnology accommodates heterogenous FAP expression in patients^31^. Additionally, intravenous administration of NPs and our use of biomaterial components found in products approved by the FDA allows for non-invasive delivery and makes our platform highly clinically translatable, overcoming complications associated with the implantation or direct injection of biomaterials to the diseased heart^33^. Lastly, our platform nanotechnology allows the exploration of a variety of therapeutic drugs, as well as other fibrotic diseases where FAP is expressed, including solid tumors (e.g. lung^48^ and pancreatic^49^ cancer) and fibrotic diseases of divergent organ-systems (e.g. liver^37,38^ and lung^39^ fibrosis), enabling broad translational impact beyond cardiovascular disease.

## Methods

### Biomaterials, drug and dyes

Chitosan (MW = 310-375 kDa, Sigma-Aldrich), sodium hyaluronate (HA, MW = 70 kDa, LifeCore Biomedical), talabostat mesylate (talabostat, MW = 310.2 Da, Cayman Chemical), sulfo-Cyanine7 carboxylic acid (Cy7, MW = 746.97 Da, Lumiprobe Corp) and fluorescein isothiocyanate–dextran (FITC-dextran, MW = 10 kDa, Sigma-Aldrich) were used as purchased. Acrylated HA was synthesized as previously described^50^. Briefly, HA was dissolved at 2 wt/ vol % in deionized water for chemical reaction with acrylic anhydride (MW = 126.11, Accela ChemBio Inc) maintained at pH 8.5-9.5 for 12 hours by 5 N sodium hydroxide (Sigma-Aldrich). Synthesized acrylated HA was purified by dialysis against deionized water and lyophilized to obtain a white powder. Esterification efficiency from dissolving acrylated HA at 1 wt/ vol % in D_2_O was calculated to be ~67% from ^1^H NMR (400 MHz Bruker) using MestReNova software (version 14.2.0-26256).

### Nanoparticle (NP) formulation and crosslinking

Calcium chloride (MW = 110.98 Da, Sigma-Aldrich) and chitosan, each at 0.5 wt/ vol %, were dissolved in 2 vol/ vol % acetic acid (MW = 64.08 Da, Thermo Fisher Scientific); where needed, talabostat (2 mg/ mL), Cy7 (0.0124 mM), or FITC-dextran (0.01 wt/ vol %) was dissolved in this solution. The chitosan solution containing calcium chloride and the desired drug or dye was added dropwise to an acrylated HA solution (1 wt/ vol %, stirred at 800-1000 at room temperature) in basic deionized water (pH 8.0) at a rate of 90 mL/hr using an injection pump. The solutions were mixed in equal volumes and then filtered through a 1 µm cell strainer (pluriSelect) to remove larger particles. The filtrate, containing submicron particles, was then purified by ultracentrifugation at 100,000 x g for 15 minutes using a Beckman L8-70M Ultracentrifuge, followed by two washes with basic deionized water (pH 8.0). The purified NPs were snap-frozen and lyophilized and the supernatants were assessed for encapsulation efficiency as previously described^26,27^. We observed that talabostat and Cy7 had encapsulation efficiencies (mean±SD) of 86.3±0.9 % (n = 3) and 78.1±1.5 % (n = 3), respectively, through assays described below (Drug and dye release studies).

NPs were crosslinked using the peptide GCNS*GP*SNCG (MW = 894.93 Da, GenScript Biotech). Thiols in cysteines crosslink acrylates by Michael Addition, and *GP* is cleavable by FAP enzyme. NPs were crosslinked by incubating 50 mg of NPs with 2.5 mL of 14.5 mM peptide in basic deionized water (pH 10.0) for 15 minutes. Following crosslinking, NPs were washed with deionized water by ultracentrifugation and then lyophilized. Uncrosslinked controls were exposed to similar conditions (basic deionized water (pH 10.0) for 15 mins), without inclusion of the peptide sequence. NPs were stored at −20 °C until used.

### Monoclonal antibody adsorption to NPs and characterization

For characterization studies, NPs were suspended in 2 vol/ vol % acetic acid at 0.21 mg/ mL. Anti-mouse FAP (Catalog #BE0374) or mouse IgG1 (Catalog #BE0083) isotype control (BioXCell) were each suspended at 0.125 mg/ mL in 10 mM HEPES buffer (pH 7.4) as previously described^51^. Zeta potential of antibodies and NPs were measured before further experiments using a Zetasizer Nano ZS with Zetasizer software (version 7.11, Malvern Panalytical). To adsorb antibodies to NPs, a 1:16 dilution (vol/ vol in final solution) of NP suspension in HEPES buffer (containing antibodies) was prepared, with a final pH of 5.0-5.5. This system (pH of ≥ 5.0 and HEPES buffer) was designed to mimic HEPES buffer used in preparing certain medications in clinics, and the pH range of sterile water for injection (US Pharmacopeia). We assessed the effect of antibody adsorption on overall NP particle charge at different wt/ wt ratios of NPs to antibodies, including 1:0.2, 1:9 and 1:18 using the Zetasizer Nano ZS. Subsequently, in-vitro and in-vivo studies were undertaken using formulations comprising NPs to antibody wt/ wt ratios of 1:9, which was reconstituted 10 minutes before intravenous administration. NPs, alone or after adsorption to antibodies, were assessed for hydrodynamic diameters and polydispersity indices using the Zetasizer Nano ZS in suspension, then imaged by transmission electron microscopy (FEI Tecnai T12 Spirit, 120kV LaB6 filament) using AMT Image Capture Engine (version 602, Advanced Microscopy Techniques) after air drying on carbon film 200 mesh Cu Grid (Electron Microscopy Sciences).

### Drug and dye release studies

Talabostat release studies (4 mg of uncrosslinked or crosslinked NPs) were undertaken in Milli-Q water (n = 3 biological replicates) at 37 °C at 60 rpm in an orbital shaker. 1mL of Milli-Q water was used to suspend NPs hourly for 6 hours, then at 1 and 2 days later to obtain releasates (supernatant), after centrifuging at 21,100 x g for 5 mins. After releasates at the 2 day timepoint were collected, NPs were incubated under similar conditions in hyaluronidase (catalog # H3884, Sigma-Aldrich) at 1 mg/ mL in 1x phosphate buffered saline (PBS; Thermo Fisher Scientific) for 48 h to measure unreleased talabostat. To measure loading content, 3.3 mg of crosslinked NPs encapsulating talabostat was suspended in 0.5 mL hyaluronidase (n = 3 biological replicates) for 48h, after which the supernatant was collected following centrifugation. We determined that 3.3 mg of NPs contained 6.2 ± 0.3 (mean±SD) ng of talabostat (n = 3) using the FAP activity assay as previously described^15^. Briefly, the Z-Gly-Pro-AMC substrate (Bachem) and recombinant human FAP enzyme were used at final well concentrations of 50 μM and 0.1 μg/ mL, respectively, in a 50 mM Tris buffer (Sigma-Aldrich) containing 1 M sodium chloride and 1 mg/ mL bovine serum albumin (Sigma-Aldrich) at pH 7.5 for the FAP activity assay. Excitation and emission wavelengths of 380 nm and 460 nm, respectively, were used to make a standard curve for talabostat on a plate reader (Infinite 200 Pro, Tecan; Tecan i-control software version 2.0.10.0).

Similarly, Cy7 release studies (2.4 mg of uncrosslinked or 4.5 mg of crosslinked NPs) were undertaken in 2 mL of Milli-Q water (n = 3 biological replicates) at 1h, 3h and 6h. For responsivity, recombinant human FAP enzyme (catalog # 3715-SE, R&D Systems) at 0.1 or 0.01 μg/ mL in Milli-Q water was used. After releasates at the 6h timepoint were collected, NPs were incubated in hyaluronidase for 48h. Cy7 release was measured based on Cy7 standards created at excitation and emission wavelengths of 740 nm and 780 nm, respectively, on a plate reader.

### Human cardiac fibroblast (HCF) culture and immunocytochemistry

Primary HCFs isolated from the left ventricles of the adult heart (PromoCell) were seeded in 96-well glass bottom plates at 10,000 per well in 200 μL complete medium as previously described^52–55^. Complete medium was made with DMEM medium, 10% heat-inactivated Fetal Bovine Serum, and 100 U/ mL penicillin–streptomycin (all from ThermoFisher Scientific). HCFs were activated using 10 ng/ mL of transforming growth factor beta-1 (TGF-β1) for 4h^56^. For in-vitro experiments assessing the effect of talabostat on FAP expression, talabostat was reconstituted in complete medium to achieve 30 nM (to mimic the peak serum concentrations of talabostat measured during clinical trials^15^), 560 nM (the half-maximal inhibitory concentration of talabostat) and 30 μM concentrations for 24h. Thereafter, we undertook immunocytochemistry (see below).

In-vitro studies were used to assess the uptake of NPs (encapsulating FITC-dextran) adsorbed with (anti-) FAP antibodies (FAP NPs) versus IgG1 controls (IgG NPs), using 6.6 μg NPs in 10 μL (mixture of HEPES buffer and acetic acid as described above) added to each well. After 24h, we undertook immunocytochemistry.

Each well was washed twice with PBS, then fixed in 10% neutral buffered formalin (Sigma-Aldrich) for 15 mins. Afterwards, 0.5% Triton X-100 (Thermo Fisher Scientific) was used for permeabilization, followed by a 30 min incubation step in 0.5% bovine serum albumin (Sigma-Aldrich). Afterwards, each well was incubated with mouse monoclonal anti-≥-SMA (1 *µ*g/ mL, catalog # MAB1420, R&D Systems) and human polyclonal anti-FAP (2 *µ*g/ mL, catalog # AF3715, R&D Systems) in PBS. After three wash steps with PBS, wells were incubated at room temperature for 30 mins with DAPI (1 *µ*g/ mL, catalog # D1306), goat anti-mouse IgG AF-568 (2 *µ*g/ mL, catalog # A-11004) and donkey anti-sheep IgG AF-647 (2 *µ*g/ mL, catalog # A-21448), all sourced from Thermo Fisher Scientific. Afterwards, wells were washed with PBS and imaged by confocal microscopy.

Fluorescent z-stacks of stained cells were taken on an Eclipse Ti2 confocal microscope (NIS Elements imaging software version 6.02.03, Nikon), equipped with a 20x water immersion objective lens. All images were analyzed by counting nuclei and applying thresholds to obtain areas using Fiji software (version 2.14.0/1.54f, NIH). Quantitative data were derived from n = 3 different wells, with 3 fields of view obtained from each well. Additionally, representative images were similarly adjusted for brightness and contrast.

### Ischemia-reperfusion model of myocardial infarction (MI)

We used an ischemia-reperfusion injury model approved by the Institutional Animal Care and Use Committee at the University of Colorado Boulder (approval number: 2871) to surgically induce MI in ten-to twelve-week-old male Wistar rats (Charles River) weighing approximately 300 g. Before surgeries, rats were weighed and, where needed, echocardiography (described in detail below) was undertaken. For preoperative analgesia, rats received a single subcutaneous injection of sustained release buprenorphine (1.2 mg/ kg). Afterwards, animals were induced in 5 % isoflurane before being intubated intratracheally for mechanical ventilation using 2 % isoflurane. After surgical scrub, a left thoracotomy at the 4th intercostal space was performed, with lidocaine infiltration in the intercostal muscles. Following the placement of retractors, the left anterior descending (LAD) coronary artery was identified and ligated using a 6-0 silk suture tied around a sterile monofilament nylon suture (LOOK 1, Hospeq Medical Equipment and Supplies). The chest retractors were removed and the skin was temporarily apposed, while the animal remained ventilated during the 30-minute ligation. Afterwards, the ligature around the LAD was removed for reperfusion, which is visualized by the return of color to the myocardial tissue. Inclusion criteria for in-vivo studies included successful ligation of the visualized LAD, confirmed by tissue blanching distal to the suture and entry of blood into the hub of the needle, respectively.

### Biodistribution and therapeutic efficacy

Alongside healthy controls, rats with MI were randomly assigned to 4 groups (n = 6 rats per group) for biodistribution studies 3 and 7 days after MI. These groups included rats with MI or healthy controls that received IgG NPs and FAP NPs encapsulating Cy7. Each rat intravenously received a 4mg NP formulation on day 3 or 7 post-MI, and was euthanized one hour later. Following euthanasia, the heart, liver, lung, spleen and kidneys were collected, snap-frozen in liquid nitrogen and stored at −80 °C for in-vivo imaging (IVIS). For IVIS imaging, organs from all four groups were imaged at once to enable side by side comparison. The IVIS Lumina III with Living Image software (version 4.8.0, Revvity) was used for both image acquisition and analyses.

To appraise therapeutic efficacy, we included sham controls (open thoracotomy but without ligation of the LAD; n = 10 rats), MI groups that intravenously received no treatment (n = 10 rats), empty NPs without drug (n = 10 rats), NPs encapsulating talabostat (n = 9 rats) or the equivalent amount of talabostat via normal saline (n = 11 rats) to mimic systemic drug administration. Each rat received 66 μg of empty or talabostat-loaded NPs, adsorbed with FAP antibodies, in a 500 μL volume by tail vein injection on day 3 and 7 after MI. Based on ascertained loading content of NPs encapsulating talabostat (3.3 mg of NPs contained 6.2 ng of talabostat), we administered the equivalent amount of talabostat present in 66 μg of NPs (0.125 ng) via 500 μL of normal saline. Data from n = 2 rats (empty NPs without drug) were excluded by Grubb’s outlier test. Body weights were taken at day 30 post-MI, and non-invasive echocardiography measurements of left ventricular (LV) dimensions and function were acquired and analyzed using a Vevo 3100 ultrasound imaging system (using software Vevo Lab version 5.6.1, Visual Sonics). Briefly, under general anesthesia (2% isoflurane), the surgical site was shaved and depilatory cream (Nair) applied before ultrasound imaging, after which rats were humanely euthanized. Whole organs, including the heart, liver, lung, spleen, kidneys and skin were collected for histological assessments.

### Tissue sectioning and infarct assessment

Following echocardiography (therapeutic efficacy studies), freshly collected heart tissues were sectioned on a rat heart slicer matrix (Supplementary Fig. 9a), where 2 mm thick sections were stained using a 1 wt/ vol % 2,3,5-triphenyltetrazolium chloride (TTC; Sigma-Aldrich) solution (made in PBS) at 37°C for 30 mins. Afterwards, sections were imaged on the SMZ18 stereoscope, before being fixed in 10% neutral buffered formalin for histological assessment. Using Fiji software to manually draw regions of interest, the infarct area was expressed as a percentage of the area of the entire left ventricle. As there were no infarcts in sham controls, no areas were drawn for this group.

Following IVIS imaging (biodistribution analysis) and echocardiography (therapeutic efficacy studies), organs (including the heart, liver, lung, spleen, kidneys and skin) were fixed in 10% neutral buffered formalin for one day. Thereafter, they were passed through increasing concentrations (15% and 30%) of sucrose (Sigma-Aldrich) daily. Subsequently, tissues were embedded in optimal cutting temperature (O.C.T.) compound (Tissue-Tek) by snap freezing. Multiple successive 10 µm sections were obtained using a microtome-cryostat for histological assessments.

### Immunohistochemistry of heart tissue

Slides with cryo-sectioned heart tissues were brought to room temperature then washed for 5 mins in PBS before histology. For immunohistochemistry (IHC), 0.5% Triton X-100 was used for permeabilization, followed by a 30 min incubation step in 0.5% bovine serum albumin. Afterwards, each slide was incubated with mouse monoclonal anti-≥-SMA (1 *µ*g/ mL, catalog # MAB1420, R&D Systems) and human polyclonal anti-FAP (2 *µ*g/ mL, catalog # AF3715, R&D Systems) in PBS. After three wash steps with PBS, slides were incubated at room temperature for 30 mins with DAPI (1 *µ*g/ mL, catalog # D1306), goat anti-mouse IgG AF-568 (2 *µ*g/ mL, catalog # A-11004) and donkey anti-sheep IgG AF-647 (2 *µ*g/ mL, catalog # A-21448). Afterwards, slides were washed with PBS, then aqueous mounting medium (Agilent) and coverslips were applied. Images of FAP expression and colocalization with administered NP formulations at day 3 and 7 post-MI were acquired on a VS200 slide scanner (software ASW 4.2, Olympus Life Science) and images of FAP expression after studies on therapeutic efficacy (30 days post-MI) were acquired by confocal microscopy. Images on therapeutic efficacy were analyzed using Fiji software by applying thresholds, where areas of FAP expression were quantified, with representative images similarly adjusted for brightness and contrast for FAP expression.

### Evaluating collagen deposition in fibrosis

For picrosirius red staining, a standard stain kit (VitroVivo Biotech) was used to stain cryo-sectioned tissues on slides according to manufacturer’s instructions. Briefly, a Weigert’s Hematoxylin working solution was used to stain the slides for 8 mins. After three wash steps in tap water, staining with picrosirius red solution for 1h was undertaken. Slides were sequentially washed in acidified water and deionized water, followed by dehydration and clearing in ethanol and xylene, respectively. Thereafter, permount mounting medium (Thermo Fisher Scientific) and coverslips were applied.

With Masson Trichrome staining, slides were incubated in pre-warmed Bouin’s Fluid (Polysciences) at 35 °C for 1h. Slides were sequentially incubated with Weigert’s Hematoxylin working solution (Polysciences), Biebrich Scarlet-Acid Fuchsin (Polysciences), phosphotungstic/phosphomolybdic acid (VWR Chemicals) and aniline blue (Polysciences), with wash steps in tap water in between each solution. Final rinse steps with acidified water were followed by dehydration and clearing in ethanol and xylene, respectively. Permount mounting medium and coverslips were applied before imaging.

Slides stained for Masson Trichrome and picrosirius red were imaged on both the SMZ18 stereoscope (NIS Elements imaging software version 5.22.00, Nikon) and the ECLIPSE Ts2 microscope (NIS Elements imaging software version 1.20.00, Nikon). Slides stained for picrosirius red were also imaged using an ECLIPSE Ci POL polarized microscope at 45° polarization angles to observe alignment (with software, Windv2 version 1.0.171206 and Instec App version 10.075P211206). Fiji software was used to manually draw regions of interest around the blue areas delineated by Masson Trichrome staining, which was expressed as a percentage of the area of the entire left ventricle. As there were no fibrotic areas in sham controls, no areas were drawn for this group.

### Histopathology

Histopathological evaluation on sectioned liver, lung, spleen, kidneys and skin was undertaken by Hematoxylin and Eosin staining. Briefly, cryo-sectioned tissues were stained in Hematoxylin staining solution (Electron Microscopy Sciences) for 30 seconds, followed by repeated wash steps in tap water and PBS. Thereafter, slides were passed through increasing concentrations of ethanol (70 and 95 %), before staining in alcoholic Eosin Y solution (Sigma-Aldrich) for 30 seconds, followed by dehydration and clearing in ethanol and xylene, respectively. Thereafter, permount mounting medium and coverslips were applied. Imaging was undertaken using the ECLIPSE Ts2 microscope.

### Statistics and reproducibility

Statistical software (GraphPad Prism Version 10.4.0 (527)) was used to analyse data, which is presented as mean and standard deviation (SD). Exact statistical test, p-values and sample sizes are provided in figure legends.

## Data availability

The data supporting the findings of this study are available within the paper and its Supplementary Information.

## Acknowledgements

This work was supported by the American Heart Association (24POST1187647 postdoctoral fellowship to C.V.M. and 25PRE1373201 predoctoral fellowship to M.O.), the American Society for Engineering Education (eFellows award to G.J.R.R), the Deutsche Forschungsgemeinschaft (DFG Project 551186431 to M.G.S.), the National Science Foundation (Graduate Research Fellowship DGE 2040434 to K.W.), and the National Institutes of Health (R01HL167994 to F.G.S. and J.A.B.). We thank the Shared Instruments Pool (SIP) Core Facility (RRID: SCR_018986), BioFrontiers Advanced Light Microscopy Core (RRID: SCR_018302 and supported by NIH 1S10RR026680-01A1), Electron Microscopy Services Core Facility in the Department of Molecular, Cellular and Developmental Biology, and Joselle McCracken and Timothy White at the University of Colorado Boulder for use of their equipment. Any opinions, findings, and conclusions or recommendations expressed in this material are those of the author(s) and do not necessarily reflect the views of funding agencies.

## Author contributions

Conceptualization, C.V.M. and J.A.B.; Methodology, C.V.M., G.J.R.R., M.G.S., A.R.P., S.B., K.W., M.O., S.R.K., F.G.S., and J.A.B.; Investigation, C.V.M., G.J.R.R., M.G.S., A.R.P., S.B., K.W., M.O., S.R.K.; Writing – Original Draft, C.V.M.; Writing – Review & Editing, C.V.M., G.J.R.R., M.G.S., A.R.P., S.B., K.W., M.O., S.R.K., F.G.S., and J.A.B.; Funding Acquisition, F.G.S., and J.A.B.; Resources, J.A.B.; Supervision, J.A.B.

## Supplementary Information

**Supplementary Figure 1.**
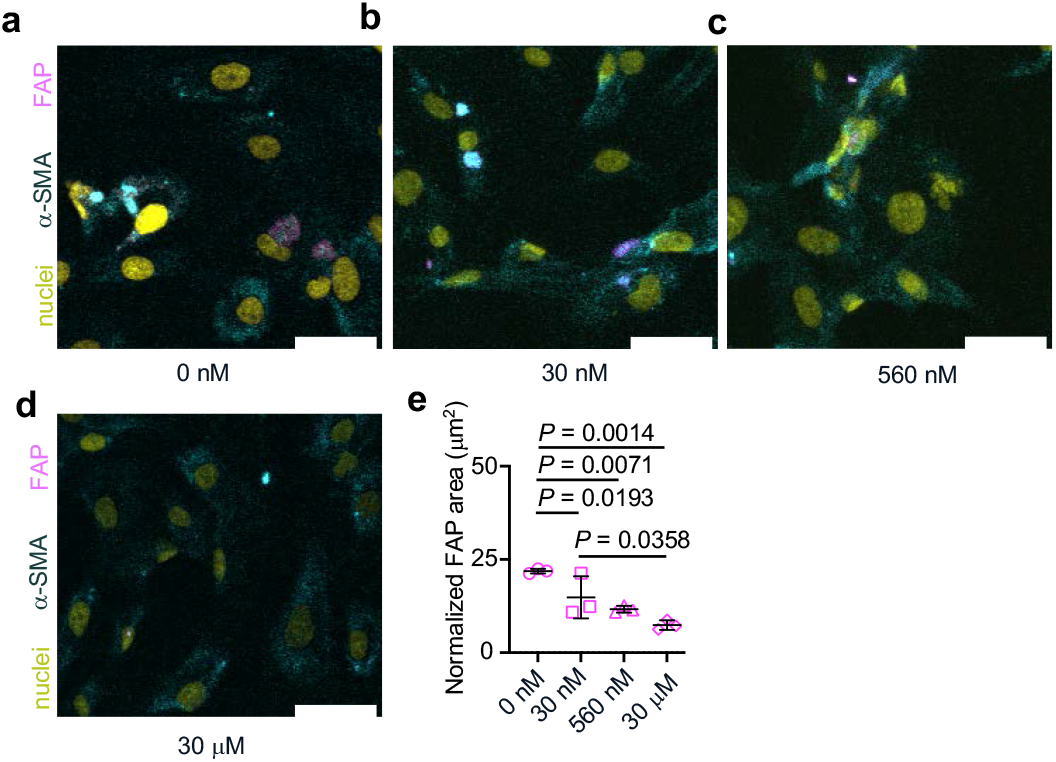
Dose-dependent effect of talabostat on FAP expression in activated human cardiac fibroblasts (HCFs). **a-d**, Immunohistochemical images of primary HCFs activated by transforming growth factor beta-1 (TGF-≥1) followed by 24h incubation with various concentrations of talabostat (scale bar, 50 μm). **e**, Quantification of FAP area normalized to cell number. Mean (SD); one-way ANOVA followed by Newman-Keuls’ multiple comparisons test; n = 3 biological replicates.

**Supplementary Figure 2.**
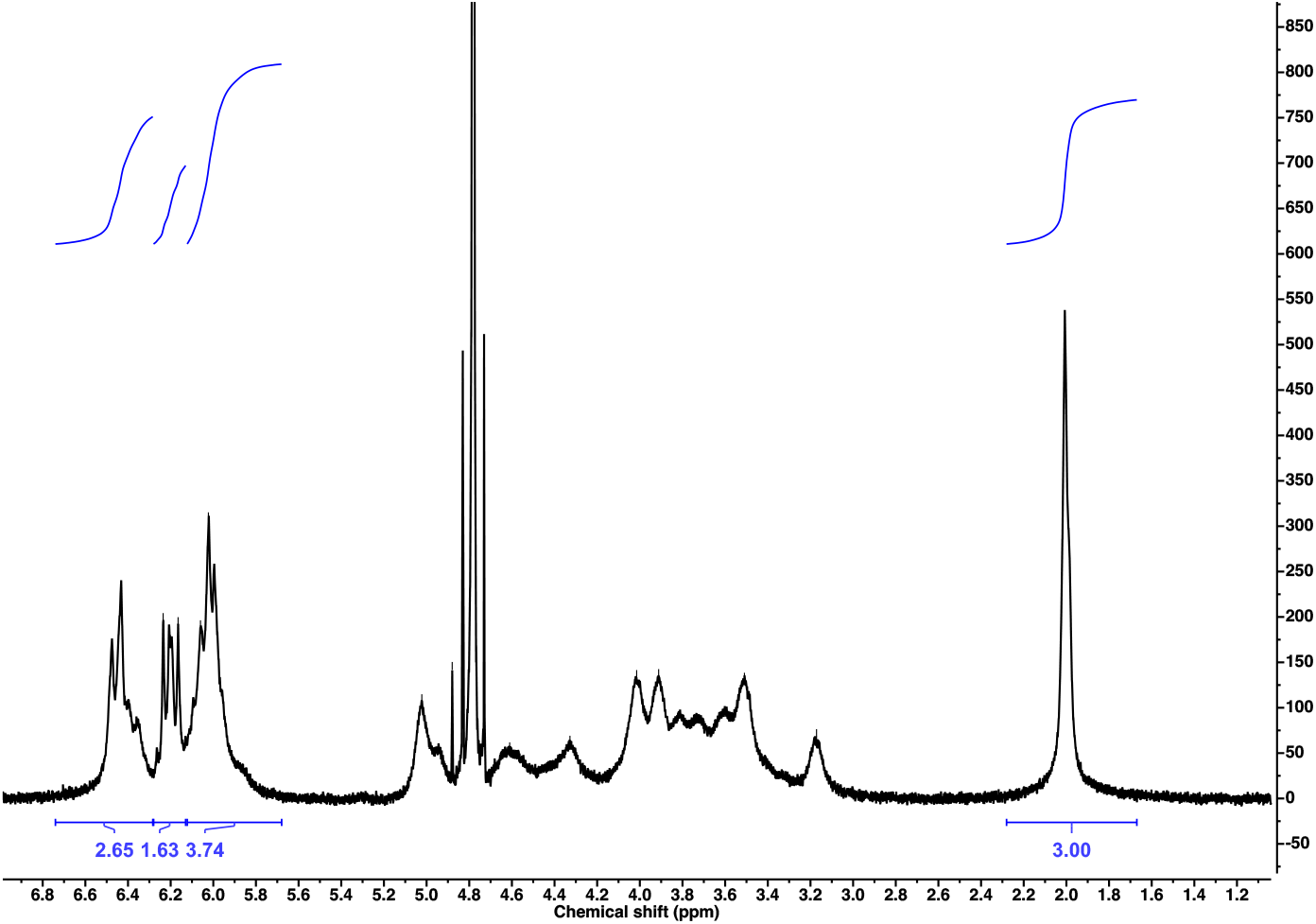
^1^H NMR spectrum of acrylated hyaluronic acid. The degree of modification determined by comparing the integrated area of the acrylate peaks to the integrated area of the methyl group on hyaluronic acid and considering all alcohol groups.

**Supplementary Figure 3.**
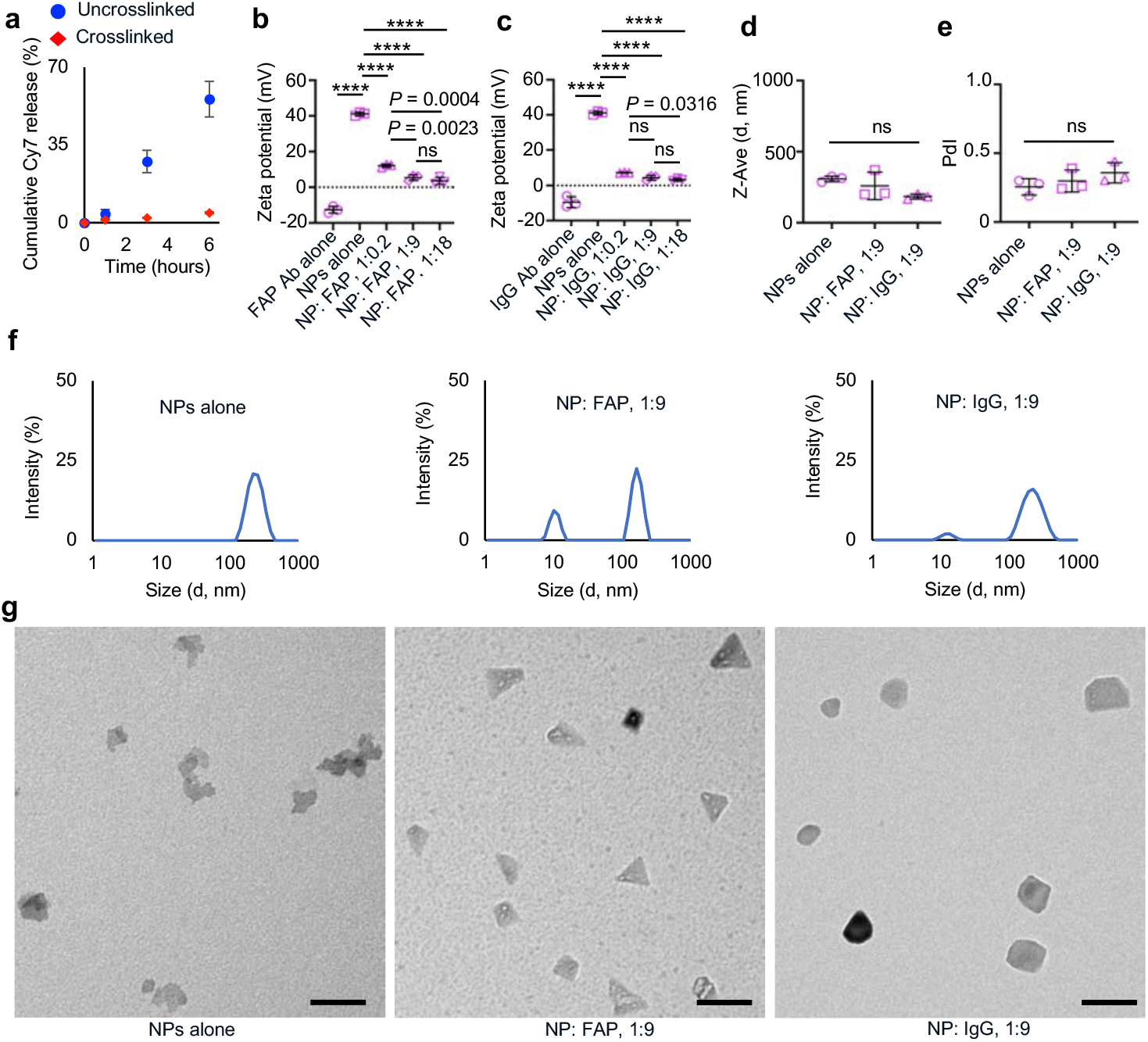
Release kinetics of Cy7 and characterization of talabostat-loaded NPs adsorbed to antibodies. **a**, Cy7 release from uncrosslinked and crosslinked NPs. **b-c**, Zeta potential of NPs, FAP and IgG antibodies (Ab), alone and in complex at various ratios. **d-f**, Hydrodynamic diameter (Z-Ave, **d**), polydispersity index (PdI, **e**) and size distribution (**f**) of hydrated NPs, alone and in complex with Ab. **g**, Transmission electron microscopy (TEM) images of dried NPs (scale bar, 100 nm). Mean (SD); one-way ANOVA followed by Tukey’s multiple comparison test (a, c, d) or one-way ANOVA followed by Newman-Keuls’ multiple comparisons test (b); n = 3 biological replicates, **** *P* < 0.0001.

**Supplementary Figure 4.**
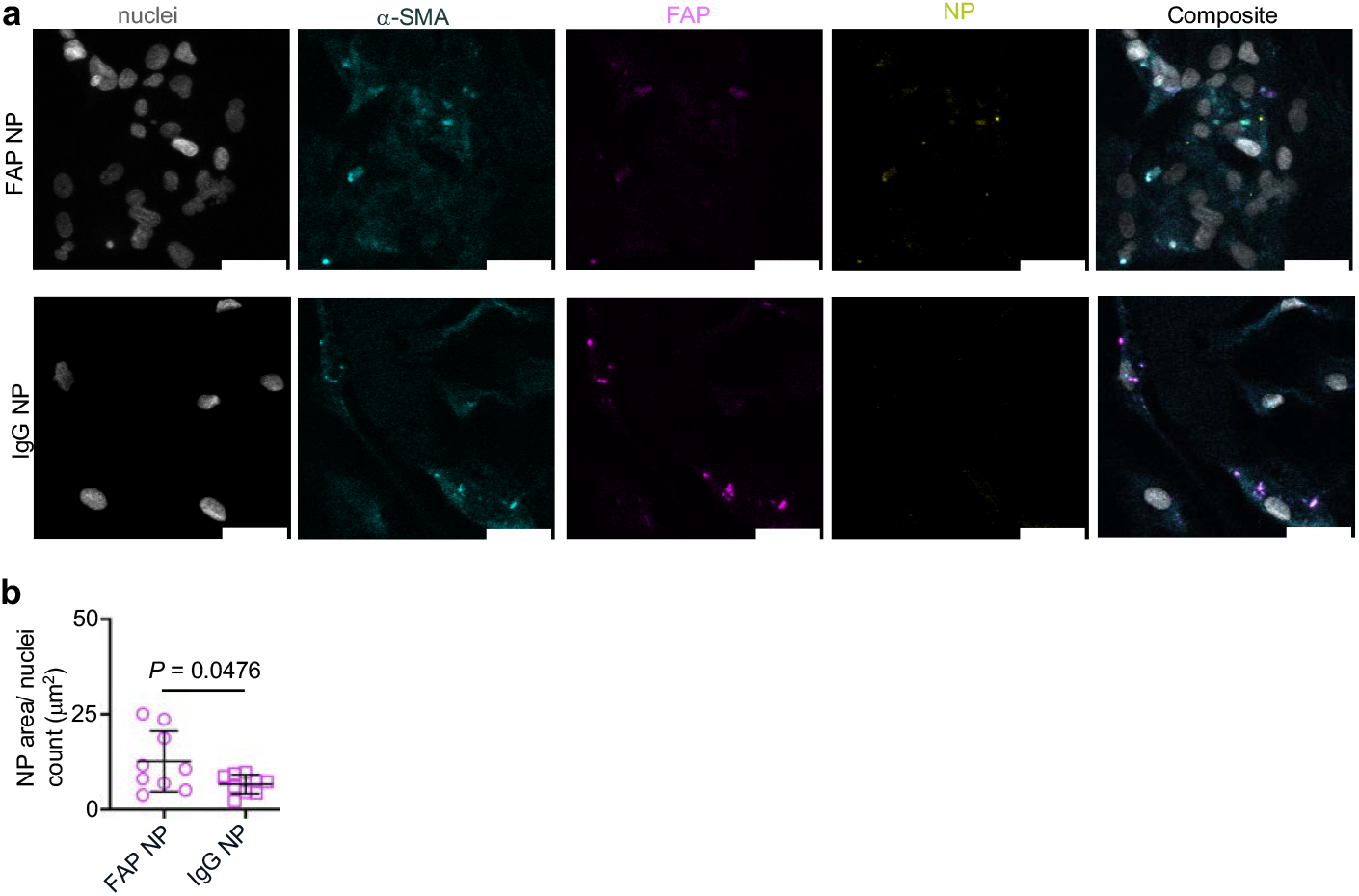
Uptake of NPs by activated human cardiac fibroblasts (HCFs). **a**, Representative immunohistochemical images of HCFs activated by transforming growth factor beta-1 (TGF-≥1) followed by 24h incubation with NPs (containing fluorescein isothiocyanate dextran) with adsorbed FAP (FAP NP; top) or IgG (IgG NP; bottom) antibodies; scale bar, 50 μm. **b**, Quantification of FAP area normalized to cell number. Mean (SD); two-tailed unpaired t-test, n = 9 fields of view obtained from 3 biological replicates.

**Supplementary Figure 5.**
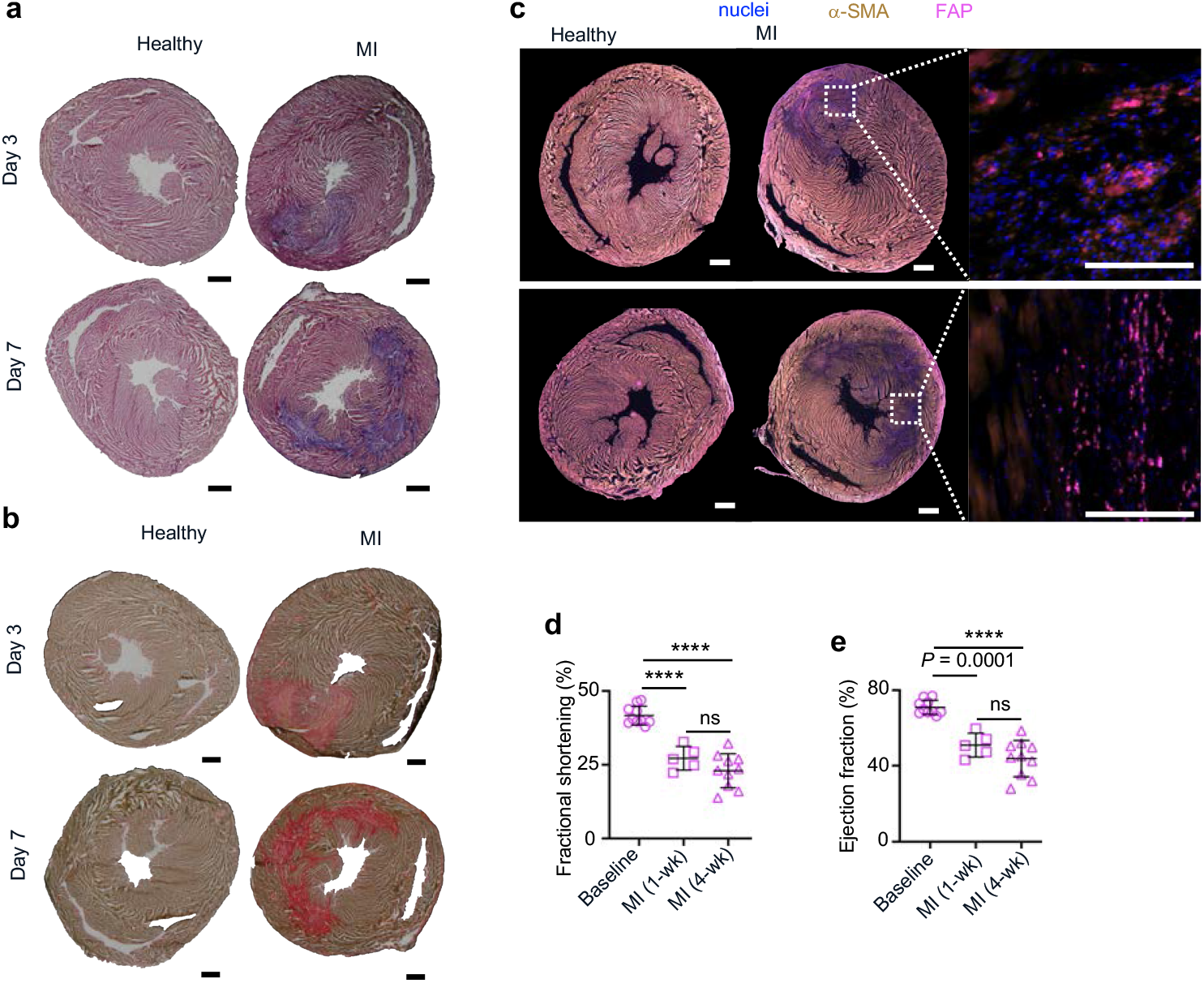
Progression of cardiac fibrosis, FAP expression, and function after MI. **a-b**, Masson Trichrome (a) and Picrosirius red staining of axial heart sections on day 3 (top) and 7 (bottom) in healthy controls and MI groups (scale bar, 1 mm). **c**, Immunohistochemical staining on day 3 (top) and 7 (bottom) in healthy controls and MI groups (scale bar, 1 mm), with magnification (scale bar, 100 μm). **d-e**, Cardiac function assessed by fractional shortening (d) and ejection fraction (e) at baseline (just before MI) and one- and four-weeks following MI. Mean (SD); one-way ANOVA followed by Tukey’s multiple comparison test; n = 10 rats (baseline), 5 rats (1-wk after MI) and 10 rats (4-wk after MI).

**Supplementary Figure 6.**
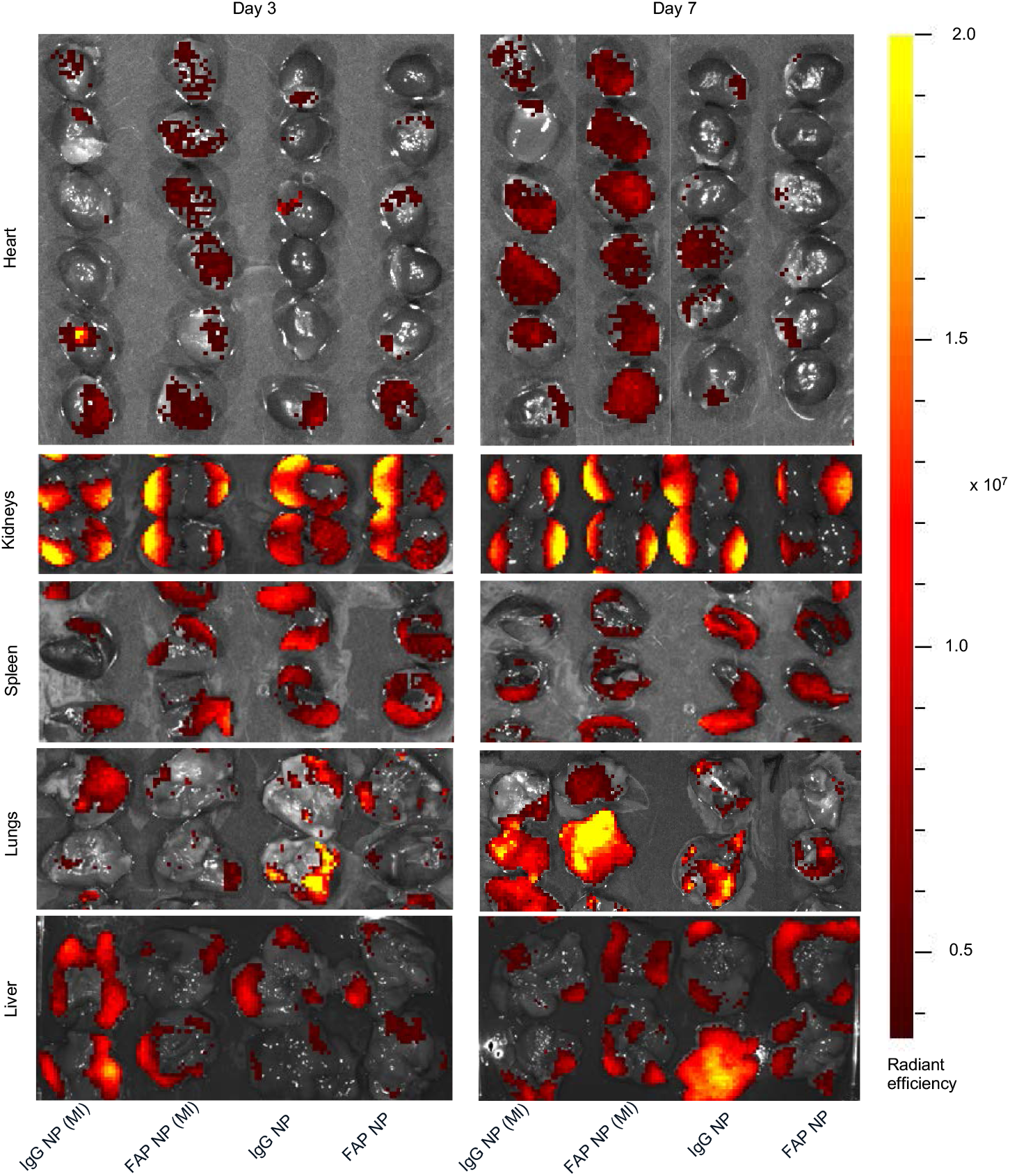
Biodistribution of administered NPs. Whole organ IVIS images obtained an hour after NPs were intravenously administered. All 6 hearts from n = 6 rats are shown, whereas kidneys, spleen, lungs and liver from n = 2 rats are shown as representative.

**Supplementary Figure 7.**
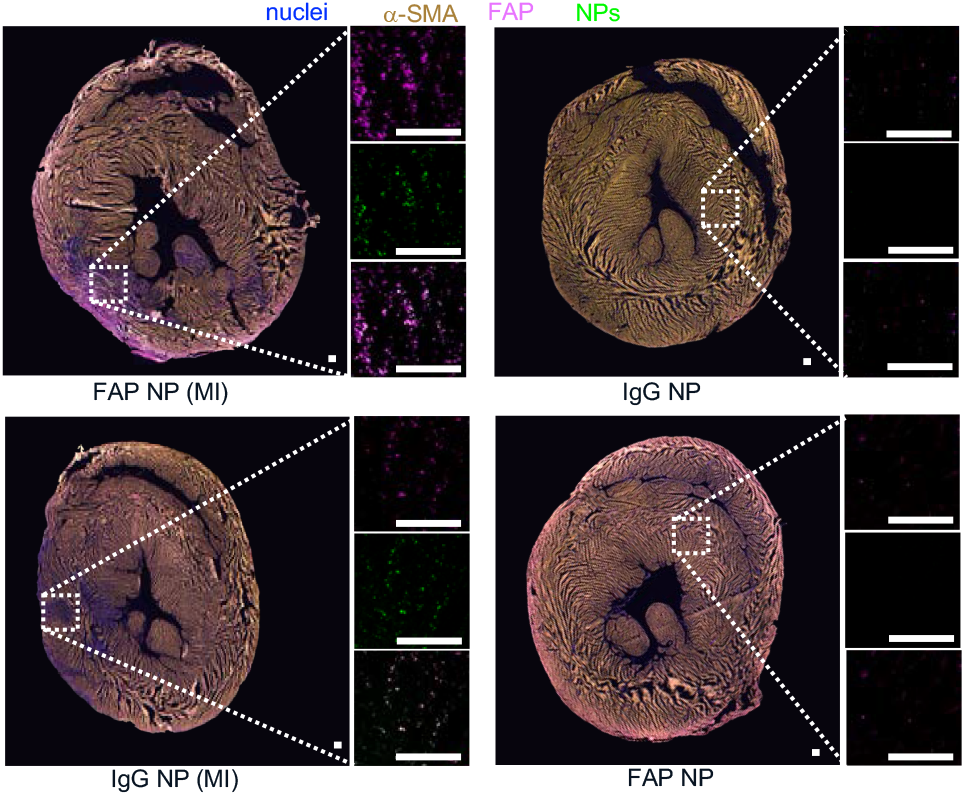
Targeting of administered NPs to the diseased region of the heart. Immunostaining (with magnification) of axial heart sections from rats with and without myocardial infarction (MI) that were injected with NPs (containing Cy7) with adsorbed FAP (FAP NP) or IgG (IgG NP) antibodies on Day 3 post-MI; scale bar, 200 μm.

**Supplementary Figure 8.**
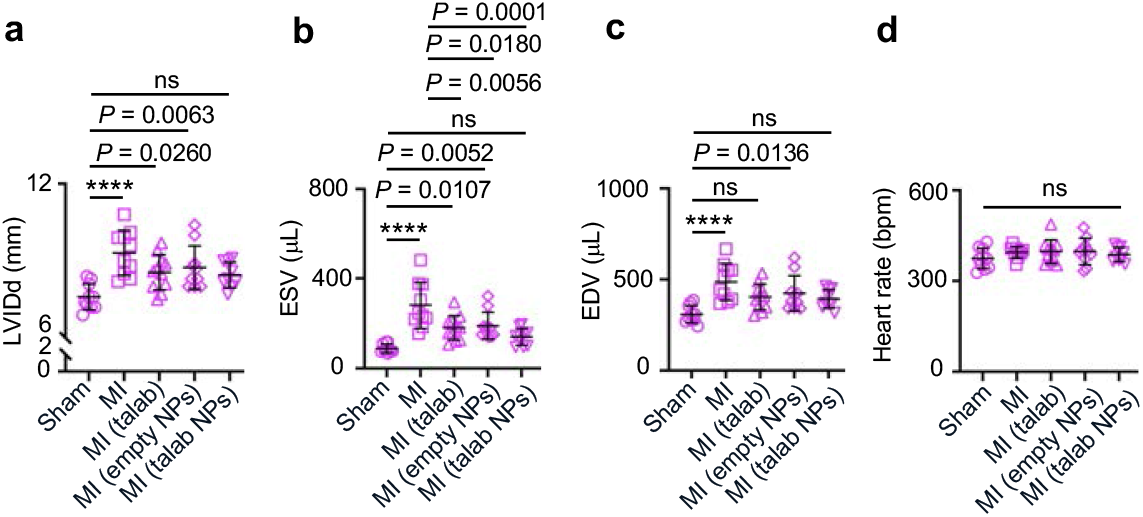
Effect of loaded NPs on additional cardiac functional indices. **a-d**, Cardiac function assessed by left ventricular internal diameter at end diastole (LVIDd; a), end-systolic volume (ESV; b), end-diastolic volume (EDV; c) and heart rate (d) for sham controls, MI alone, and MI with intravenous talabostat in saline (talab), empty NPs or NPs encapsulating talabostat (talab NPs). Mean (SD); one-way ANOVA followed by Tukey’s multiple comparison test; n = 10 rats per group (sham, MI, empty NPs), n = 11 (talab), n = 9 (talab NPs); not significant (ns), **** *P* < 0.0001.

**Supplementary Figure 9.**
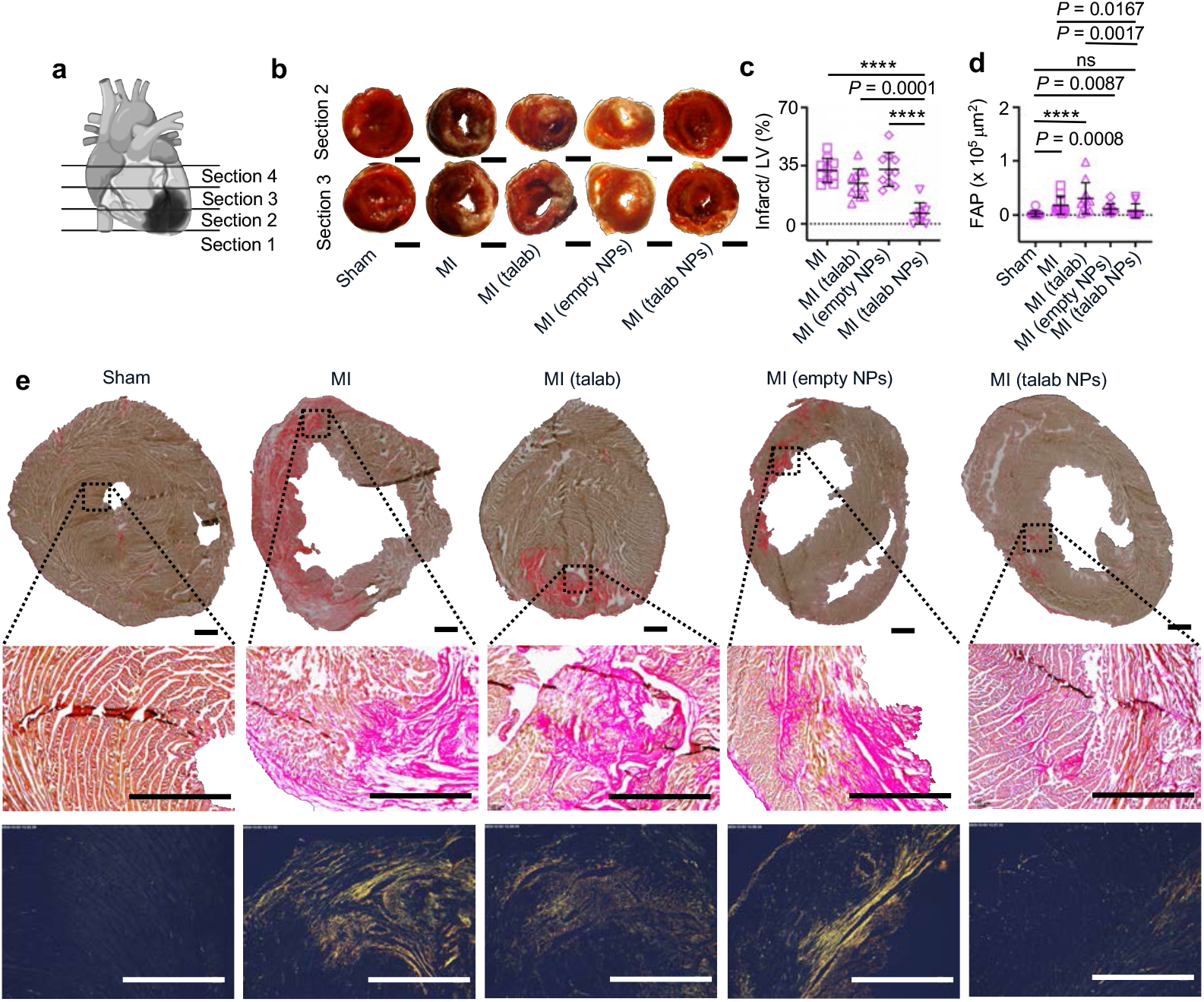
Infarct size, FAP expression, and cardiac fibrosis with and without treatment. **a**, Diagrammatic representation of heart sectioning. **b**, 2,3,5-Triphenyltetrazolium chloride (TTC) staining of heart axial sections of rats that underwent sham surgery, MI alone, MI with intravenous talabostat in saline (talab), empty NPs or NPs encapsulating talabostat (talab NPs); scale bar 5 mm. **c**, Quantification of LV tissue viability. **d**, Quantification of FAP expression in the LV. **e**, Picrosirius red staining of heart axial sections (top), with magnification (middle) under regular and polarized (bottom) light microscopy (scale bars, 1 mm). Mean (SD); one-way ANOVA followed by Tukey’s multiple comparison test (Supplementary Fig. 10c) or Kruskal-Wallis test followed by uncorrected Dunn’s multiple comparisons test (Supplementary Fig. 10d); n = 10 rats per group (sham, MI, empty NPs), n = 11 (talab), n = 9 (talab NPs); **** *P* < 0.0001.

**Supplementary Figure 10.**
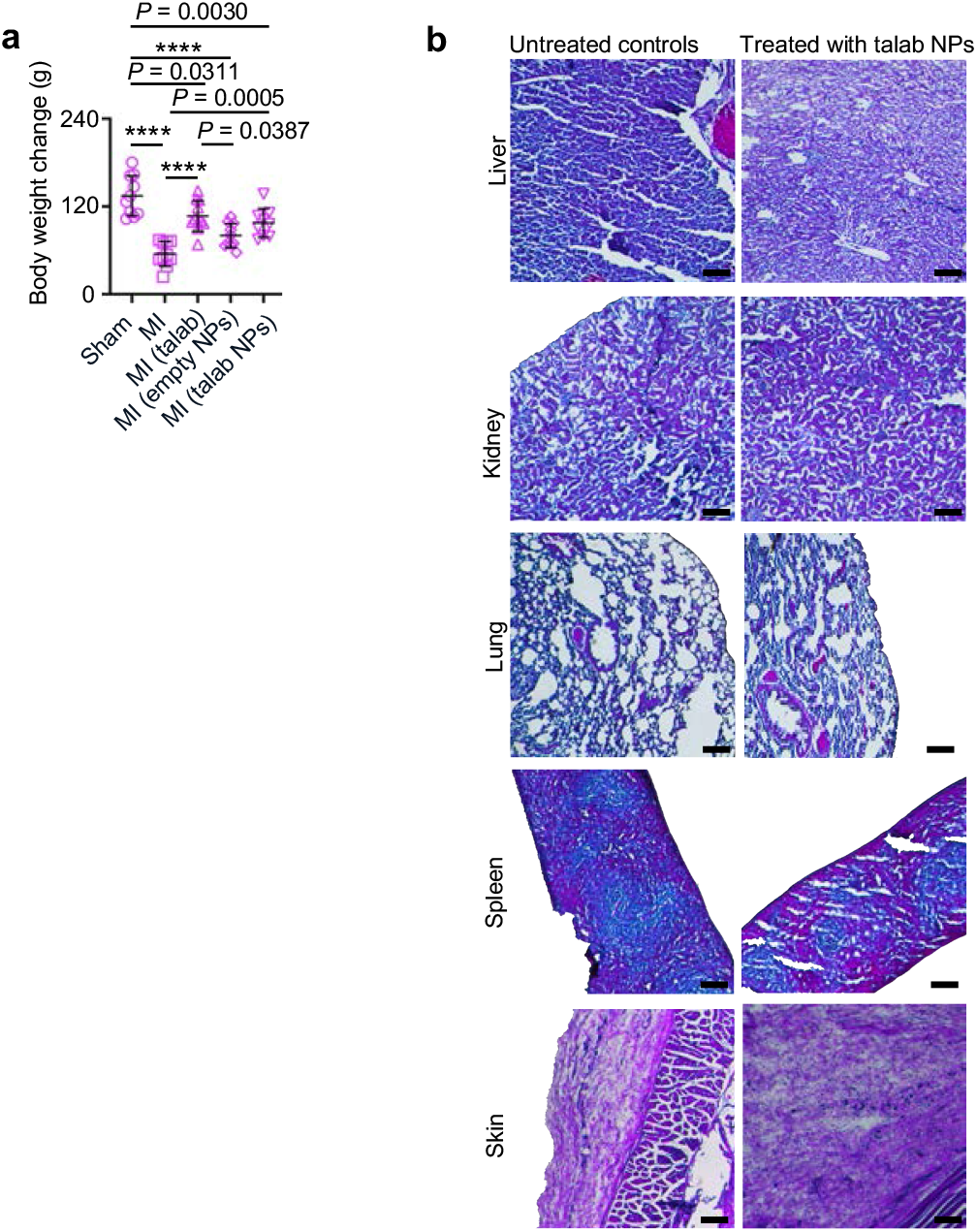
Body weight and histology of different organs following NP administration. **a**, Body weight change from baseline (before MI or sham surgery) to euthanasia after 4 weeks for sham controls, MI alone, and MI with intravenous talabostat in saline (talab), empty NPs or NPs encapsulating talabostat (talab NPs). **b**, Hematoxylin and Eosin staining of different organs of rats that received talab NPs, alongside controls that were untreated; scale bar, 200 μm. Mean (SD); one-way ANOVA followed by Tukey’s multiple comparison test; n = 10 rats per group (sham, MI, empty NPs), n = 11 (talab), n = 9 (talab NPs); **** *P* < 0.0001.

